# PIMENTO: A PrIMEr infereNce TOolkit to facilitate large-scale calling of amplicon sequence variants

**DOI:** 10.1101/2025.07.04.663168

**Authors:** Christian Atallah, Lorna Richardson, Martin Beracochea, Robert D. Finn

**Affiliations:** European Molecular Biology Laboratory, European Bioinformatics Institute (EMBL-EBI), Wellcome Genome Campus, Hinxton, Cambridge, UK

## Abstract

The identification of amplicon sequence variants from DNA metabarcoding data is a common method for revealing the taxonomic makeup of environmental samples, and for allowing comparative studies between similar datasets. A significant hurdle to the large-scale calling of amplicon sequence variants from publicly available nucleotide datasets is the heterogeneous presence of primer sequences in reads, the removal of which is a necessary pre-processing step for this form of analysis. Furthermore, as the details of the experimental primers are rarely captured in the metadata associated with the sequence records, there is a need for a method that can automatically infer the presence and identity of primers in sequencing data. In this work, we introduce PIMENTO, a Python package which uses a dual-strategy approach for identifying primers that are present in sequencing reads to enable their removal, and therefore facilitate amplicon sequence variant calling at scale.

## 1. Introduction

DNA metabarcoding is a molecular technique that targets a diagnostic marker gene and obtains genetic sequences that can be assigned to a taxonomy. This method is typically applied to genetic material present in environmental samples termed environmental DNA (eDNA), which is increasingly being applied to elucidate the organisms present in an ecosystem**[1]**. Depending on the taxonomic group of interest in a metabarcoding experiment, there are different marker genes that are typically chosen for amplification and DNA sequencing. For example, the mitochondrial cytochrome C oxidase subunit I (COI) gene is commonly used to resolve eDNA from macro-organisms like fish and insects, while the ribosomal RNA (rRNA) gene is a typical marker used for microbial diversity resolution, including bacteria and archaea, but even some eukaryotes like fungi and plankton. The use of metabarcoding experiments to reveal the taxonomic composition of samples is popular due to cost-effectiveness, standard protocols and ease of use.

One common approach for the analysis of metabarcoding sequence data is the clustering of reads into high sequence identity groups, termed operational taxonomic units (OTUs). These OTUs can then be taxonomically annotated by comparing to marker-gene-specific databases (Table 1), a process termed closed-reference OTU picking. Closed-reference OTUs therefore help describe a sample’s taxonomic diversity, and can be compared across samples when used with a consistent reference database. However, the OTU clustering can easily hide diversity at the lowest taxonomic ranks like species, including unknown organisms that might be of interest. Another approach for the analysis of metabarcoding data, which is growing in popularity, is the calling of amplicon sequence variants (ASVs). ASVs have various benefits compared to OTUs, including more precise resolution down to single-nucleotide differences, and more powerful cross-study comparisons**[8]**, though comparisons can strictly only be made between experiments amplifying the same marker, utilizing a common informatics pipeline.

**Table 1:**
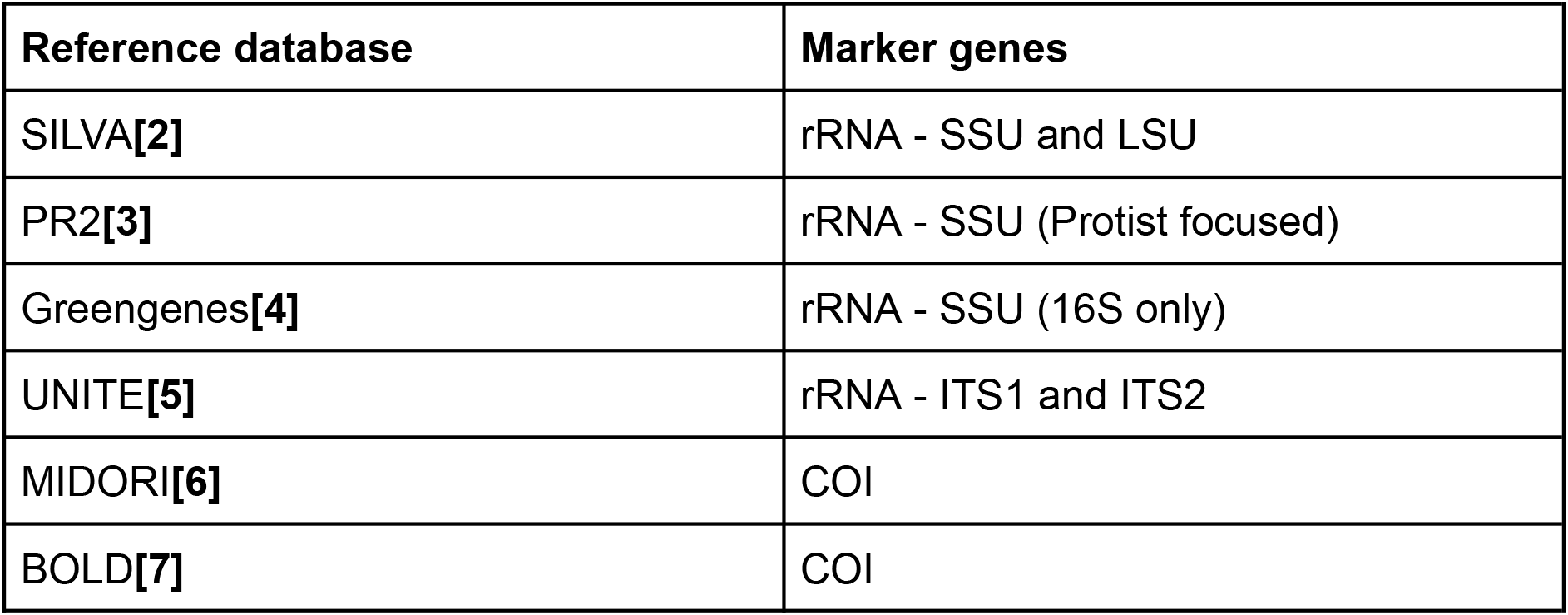
Reference databases commonly used in metabarcoding analysis, separated by marker gene.

Like many bioinformatics methods that handle sequencing reads, ASV calling necessitates a degree of quality control (QC) on the reads. Apart from common read QC steps like quality filtering and adapter trimming, the most widely used tools for generating ASVs have an additional strict requirement: the removal of synthetic nucleotides, particularly those coming from amplification primers, including DADA2**[9]** and deblur**[10]**. Synthetic nucleotides, especially ambiguous bases of primers, are likely to result in high numbers of chimeric ASVs being called, which are made up of two or more abundant “parent” ASVs. As chimeric ASVs are typically filtered out in downstream analyses, it is possible to lose the majority of the data to this filtering step if primer sequences are included in the initial ASV calling, making precise primer trimming necessary for a robust ASV analysis.

Performing primer removal on an individual study basis is usually trivial: the chosen primer sequences are normally known to the scientist conducting the experiment, which are passed to tools like Cutadapt**[11]** for primer removal. However, in larger-scale meta-analysis, where multiple studies may be considered in a single analysis, or where publicly-available data from heterogeneous sources is being analysed, the knowledge of the primer sequences used can become more variable: individual studies might use multiple different primer pairs, primers may or may not have been removed from sequence reads**[12]**. Also, within sequence records in databases like the European Nucleotide Archive (ENA)**[13]**, the experimental metadata attached to a sequence record often does not contain information about used primers**[12]**. Such variation in data availability is problematic for analysis services such as that provided by MGnify**[14]**, where automation of analysis pipelines is necessary. In this case applying the ASV analysis to vast collections of diverse data that are deposited in sequence archives such as ENA, from large studies (like Tara Oceans**[15]** and the Earth Microbiome Project**[16]**) to smaller but more specific studies, can prove more complicated.

Consequently, there is a need for a method to perform automated inference of primer sequences for the purpose of large-scale ASV calling. To this end, we introduce a tool called PIMENTO that implements a dual-strategy method for addressing this important need: it first searches a set of amplicon reads for a predefined list of standard primers that are commonly used in metabarcoding experiments, and if a match is not found a *de novo* prediction for the presence of primers not in the list is performed.

## 2. Method

PIMENTO is a Python command-line tool for performing primer inference on sequencing read files. The source code is available on GitHub**[17]**(https://github.com/EBI-Metagenomics/PIMENTO), and is packaged and released on both PyPi and bioconda for ease of distribution. PIMENTO’s dual primer-search strategy (Figure 1) is as follows:

**Figure 1:**
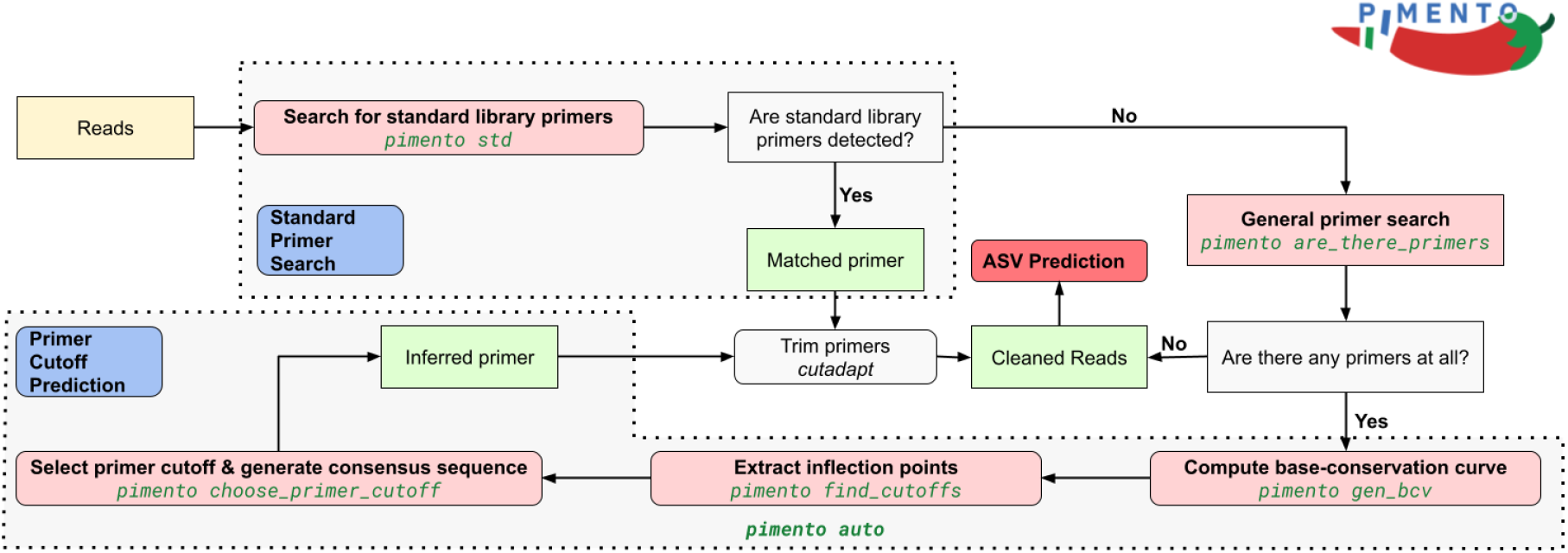
Schematic workflow of the PIMENTO tool. PIMENTO employs a dual primer-search strategy, with the standard library primer search method used first. If primers are found, they can be trimmed from the input reads using a tool like Cutadapt**[11]**. If no primers are matched, the general primer search detects if potential primers are present, then the primer cutoff prediction method is used to determine a consensus sequence by inferring a cutoff point for the primer to be trimmed.

- <u>Standard primer search:</u> fuzzy regular expression search queries to a library of curated standard primer sequences.
- <u>Primer cutoff prediction:</u> identification of the primer cutoff point from analysis of patterns of base conservation at the beginning (and end, for single-end libraries) of reads.

Below, we explain these two strategies in detail.

### 2.1 Standard primer search strategy

This first strategy utilises a manually-curated library of standard primers for metabarcoding of the ribosomal RNA (rRNA) gene. Typically, specific regions of the rRNA gene are targeted for amplification, including 16S rRNA for both bacteria and archaea, and 18S rRNA and the internal transcribed spacer (ITS1 and ITS2) for eukarya. This library was curated from a survey of the literature, sequencing platform websites, and other resources. The library currently contains 110 primers (names and sequences), of which 55 are forward strand direction (5′ to 3′) and 55 are reverse strand direction (3′ to 5′).

The strategy takes every primer sequence as a query, and performs a search against the first 50 bases of each read, the region where untrimmed primers are expected to reside. The search is fuzzy, allowing for at most two mismatches, to model for sequencing errors, and uses regular expressions, to model for ambiguous bases in some primers. The proportion of reads in which a primer is found is recorded, and any primers with a read proportion lower than a default threshold of 0.60 are disregarded. Once all primers have been searched against the reads, the primer pair with the highest proportion is identified. The default parameters for this step are optimised for the identification of a single pair of primers, but the user can modify this threshold to find multiple sets. However, changing this threshold does not assert the primer pairing, and as the threshold is lowered, the users should take more care in verifying the results.

<u>Corresponding</u> **<u>PIMENTO</u>** <u>command:</u> `pimento std <args>`

### 2.2 Primer cutoff prediction strategy

The second strategy is based on the analysis of base-conservation patterns at the start of reads to make a prediction on the “cutoff point” of the primer, i.e. where the primer sequence ends and the biological sequence begins. Specifically, it is expected that bases in the “primer space” will have high levels of conservation, as metabarcoding primers are chosen for sequence conservation by design.

Complementarily, bases in the immediate “post-primer space” are typically less conserved, as the natural sequence diversity of the organisms in the sample starts revealing itself. However, identifying the cutoff point between the “primer space” and “post-primer space” is complex, as the bases immediately after a primer can still lie in the conserved regions of amplicons like the rRNA gene. Another difficulty is caused by ambiguous bases near the end of a primer sequence, as the patterns of conservation can look similar to a true cutoff point. However, application of the method has demonstrated that the heuristic does select cutoff points that are effective in trimming reads of existing primer sequences.

This approach has one optional but recommended step, and three required steps, all of which use a common concept of “base-conservation”, which we define here. First, a window starting at the *i*-th base of the sequence and of size *n* bases is defined, the value of which will vary depending on the step of the method. Second, at each index *j* of this window, the proportion *p* of the most commonly-occurring base between A/C/T/G is computed. Finally, the mean base proportion of all the indices of the window is calculated. This mean value will represent the base-conservation *BC* starting at base *i* of the window of size *n* (Equation 1).

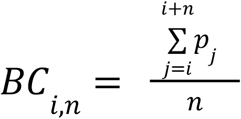

**Equation 1:** The base-conservation *BC* of the window starting at base *i* and of size *n* is defined as the mean proportion *p* of the most commonly-occurring base at each index *j*.

### Optional preliminary step

Before running the primer cutoff prediction method, it is optional but recommended to use one of PIMENTO’s utilities to flag as true or false whether there are primers present in the reads. Using base-conservation patterns, this utility splits the first 100 bases of reads into 10 different base-conservation windows of size 10 bases. By comparing the base-conservation of the first two base-pair windows, to the other eight windows, the tool will declare a primer as existing if the bases in those initial two windows are more conserved, based on the sample-specific “average” conservation across all windows.

<u>Corresponding</u> **<u>PIMENTO</u>** <u>command:</u> `pimento are_there_primers <args>`

#### Step 1

The first step of the primer cutoff prediction method slides a base-conservation window of size 5 across the first 25 indices. The sequence of base-conservation values computed by sliding this window represents a “base-conservation vector” (BCV) for the first 25 bases of the reads.

<u>Corresponding</u> **<u>PIMENTO</u>** <u>command:</u> `pimento generate_bcv <args>`

#### Step 2

In the second step, the negative inflection points of this BCV are computed. Every identified negative inflection point, i.e. a point at which the base conservation drops, is considered a potential primer cutoff point that warrants further investigation in the next step.

<u>Corresponding</u> **<u>PIMENTO</u>** <u>command:</u> `pimento find_cutoffs <args>`

#### Step 3

In the third and final step, the “best” cutoff point is chosen out of the original set of inflection points. This decision is made based on the difference between base-conservation windows before and after a point - the inflection point that has the largest difference is greedily chosen as the primer cutoff point. As a final output of this process, a consensus sequence is generated based on the previously selected cutoff point. This consensus sequence can then be used downstream to remove the primer from the reads with a tool like Cutadapt.

<u>Corresponding</u> **<u>PIMENTO</u>** <u>command:</u> `pimento choose_primer_cutoffs <args>`

For ease of use, it is possible to run the three steps of the primer cutoff inference strategy in just one PIMENTO command: `pimento auto <args>`

## 3. Results and discussion

To assess the primer inference capabilities of PIMENTO, 855 paired-end, marine microbiome, amplicon sequencing runs were randomly selected and retrieved from the ENA, comprising what we will call the Marine Survey Dataset (MSD) hereafter. The MSD was built by first querying the ENA Browser API with the following query:

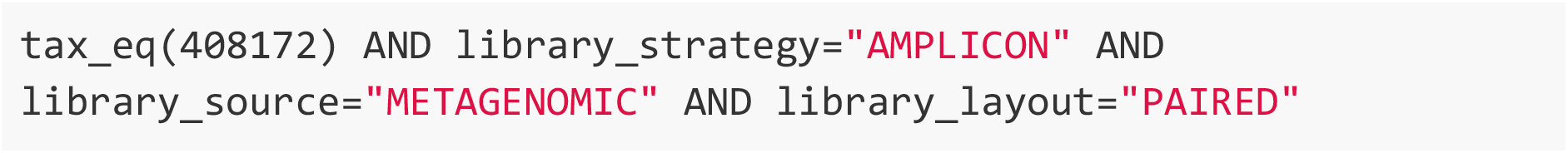

From the result of this query, 855 different studies were randomly chosen, and a random paired-end sequencing run was selected from each study. Thus, the MSD is composed of the 855 pairs of FASTQ files of these selected runs. The different steps of both the PIMENTO *standard primer search* and *primer cutoff prediction* strategies were then run on the MSD, generating primer inference results for all runs.

As popular choices of primers in metabarcoding experiments depend on the biome being studied, two additional, but smaller biome-specific datasets were built and tested the same way to assess PIMENTO’s consistency across different sampling environments. Specifically, a soil survey dataset (SSD) containing 79 random runs was built from ENA, swapping the “tax_eq” value for “410658”. Finally, a human gut survey dataset (HGSD) containing 78 runs was built from ENA, swapping the “tax_eq” value for “408170”.

### 3.1 Primer inference outcomes

The results of the *standard primer search* strategy and the optional preliminary step of the *primer cutoff prediction* strategy for all runs of the MSD are summarised in Figure 2. For both forward and reverse FASTQ files, flags were assigned depending on the outcome of each strategy.

**Figure 2:**
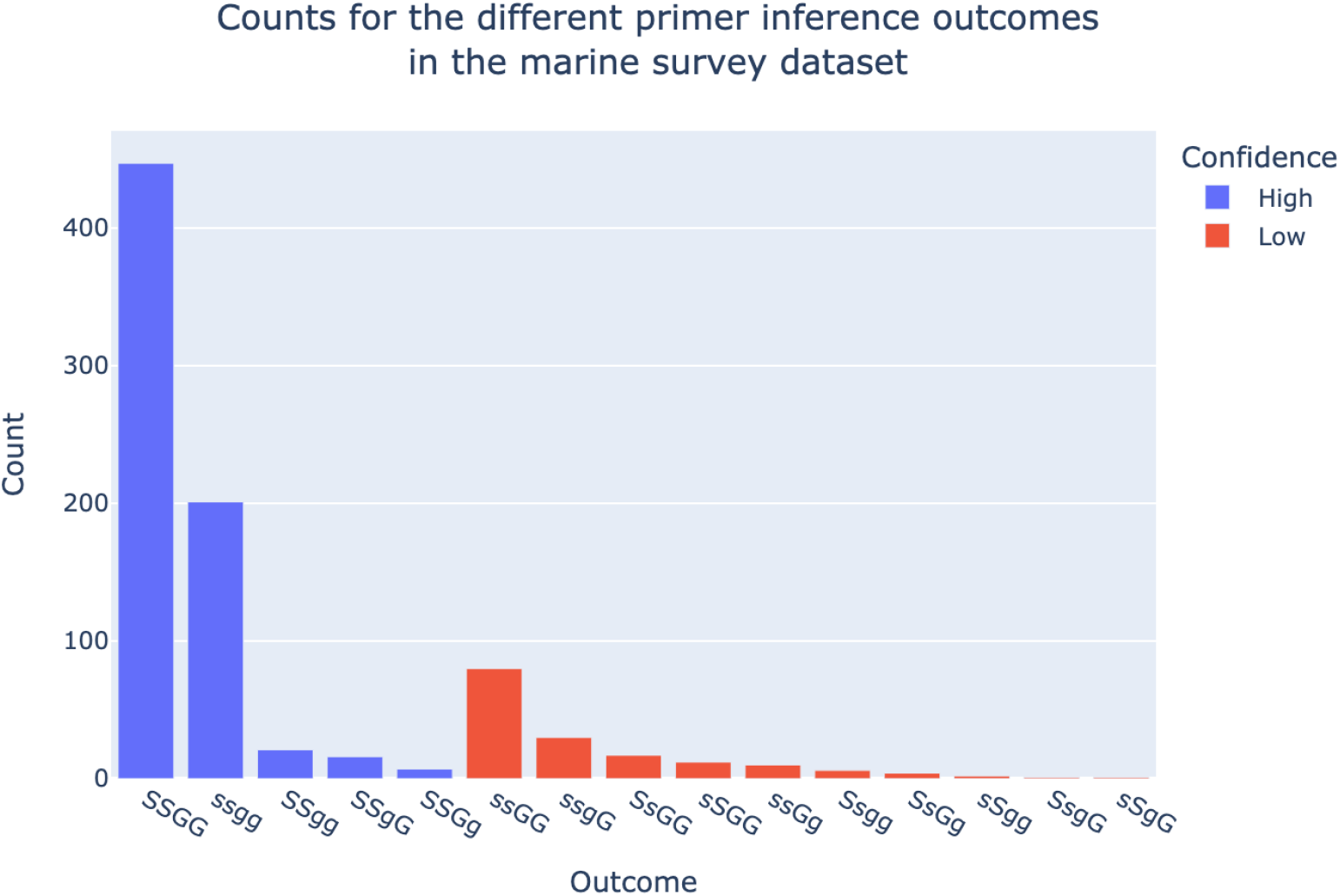
Bar plot showing the distribution of primer inference outcomes for the 855 runs of the Marine Survey Dataset. The outcomes are classified into high (blue) and low (red) confidence depending on the likely accuracy of the outcome category, with 80.9% of the outcomes considered high confidence.

- “S” - if a standard primer was found in a direction
- “s” - if no standard primer was found in a direction
- “G” - if a primer was predicted by the cutoff prediction in a direction
- “g” - if no primer was predicted by the cutoff prediction in a direction

For example, if the presence of primers was found to be likely in both directions by step 1 of the *primer cutoff prediction* strategy, and standard primers were found in both directions, the flag “SSGG” was assigned. If only standard primers were found, but the pipeline could not detect the presence of a primer the flag would be “SSgg”.

We assessed the different combinations of flags as high and low confidence of being accurate about the presence of primers based on the following interpretations. As the *standard primer search* strategy is more reliable, outcomes where standard primers were found on both strands were considered high confidence. Cases where neither strategies found primers (i.e. ssgg) were also considered high confidence outcomes as both methods indicated a negative presence of primers. All remaining outcomes are considered to be low confidence, as they mostly include cases with negative standard primer presence in combinations that are harder to interpret.

As can be seen in Figure 2, of the 855 runs, 692 were assessed to be high confidence outcomes (80.9%). This number includes 447 runs which were unanimously found to have primers by both methods (i.e. SSGG), while 201 runs (23%) have no primers (i.e. ssgg), indicating that the primers have been removed. Such a low proportion of the runs being confidently primer-free illustrates the problem and shows the current necessity for a tool like PIMENTO.

Low confidence outcomes make up 18.9% of the runs (162 out of 855), with almost half of those (80 out of 162) being from runs where the presence of primers was deemed likely in both directions, but with no matches to standard primers (i.e. ssGG). There are multiple interpretations possible for these cases: the first one - and probably the most likely - is that primers do indeed exist but are not in the standard primer library. For this scenario, the cutoff prediction method will predict an inferred primer to be trimmed, which the method does with high accuracy (see examples in section 3.3). Another possibility is that some runs will be false positives i.e. they have been flagged for primer inference even though there are no primers, which is not implausible due to the use of a heuristic method. Strategies for assessing potential false positives through primer validation are discussed in section 3.5.

### 3.2 Standard primer inferences

The results in this section pertain to runs with outcomes that were positive for standard primers in both directions (i.e every SS outcome), which total 491 runs. In Figure 3, the distributions of primer presence proportions, in both forward and reverse directions, can be seen. These proportions show that some runs had matching primers as low as 0.61, which still require trimming despite being lower value outliers. Figure 3 also shows the fraction of reads the matched primers were found in, with the first quartiles computing to 0.98 and 0.97, for the forward and reverse directions, respectively. These high values show that for the majority of runs, matched standard primers were present in the vast majority of reads, which, after trimming, would lead to cleaner reads for ASV calling purposes.

**Figure 3:**
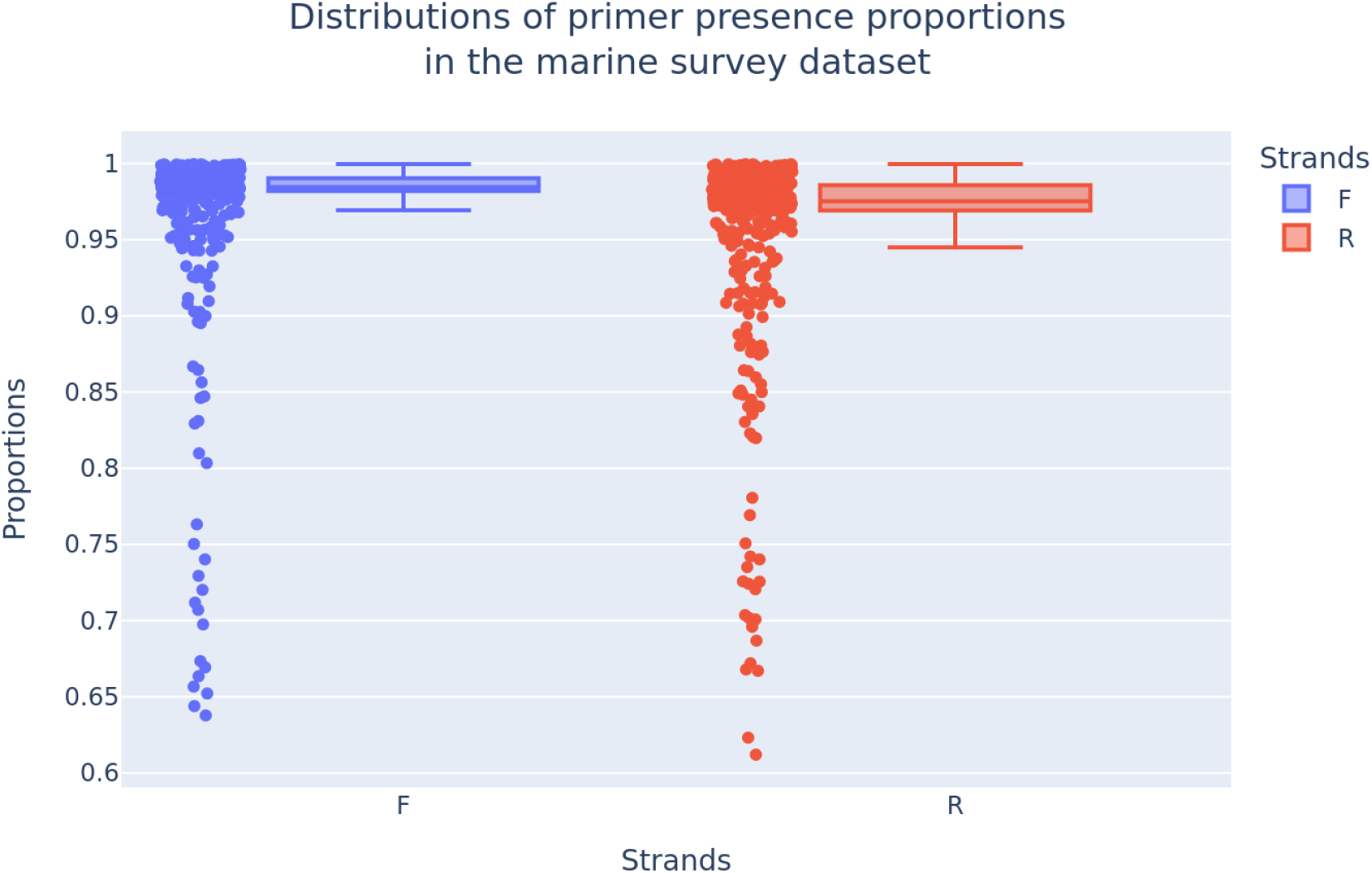
Distribution of the primer presence proportions in the Marine Survey Dataset for matched standard primers for both 5^′^ to 3^′^ (F) and 3^′^ to 5^′^ (R) directions. The vast majority of the runs had primer proportions in the reads as high as 0.97, with a minority of outliers reaching a proportion as low as 0.61.

Figure 4 plots the counts for different amplicon regions that were identified by proxy of their standard primer identity. The most common regions are 16S hypervariable regions, which is expected as the most common form of amplicon sequencing. However, the plot also shows the breadth of regions covered by the standard primer library, with occurrences of 18S, ITS, and even Archaea-specific 16S. Figure 5 shows the actual matched primers rather than the regions they amplify. The most common primers tend to be for 16S amplification, but there is a wide range of primers being represented in the MSD.

**Figure 4:**
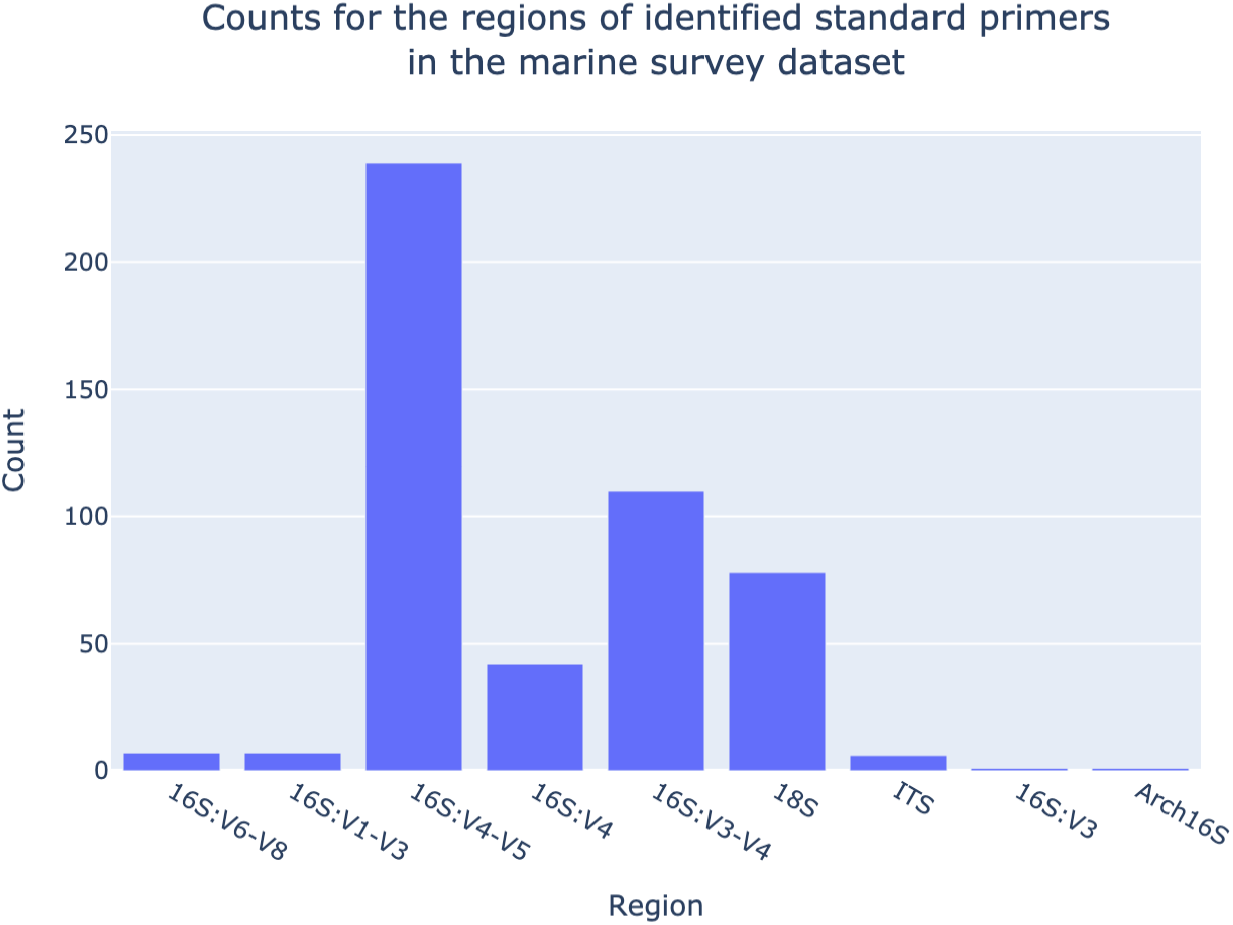
Distribution of identified amplified regions in the Marine Survey Dataset. (A) The most common amplified regions were 16S, but matches to other regions like 18S and ITS were also present.

**Figure 5:**
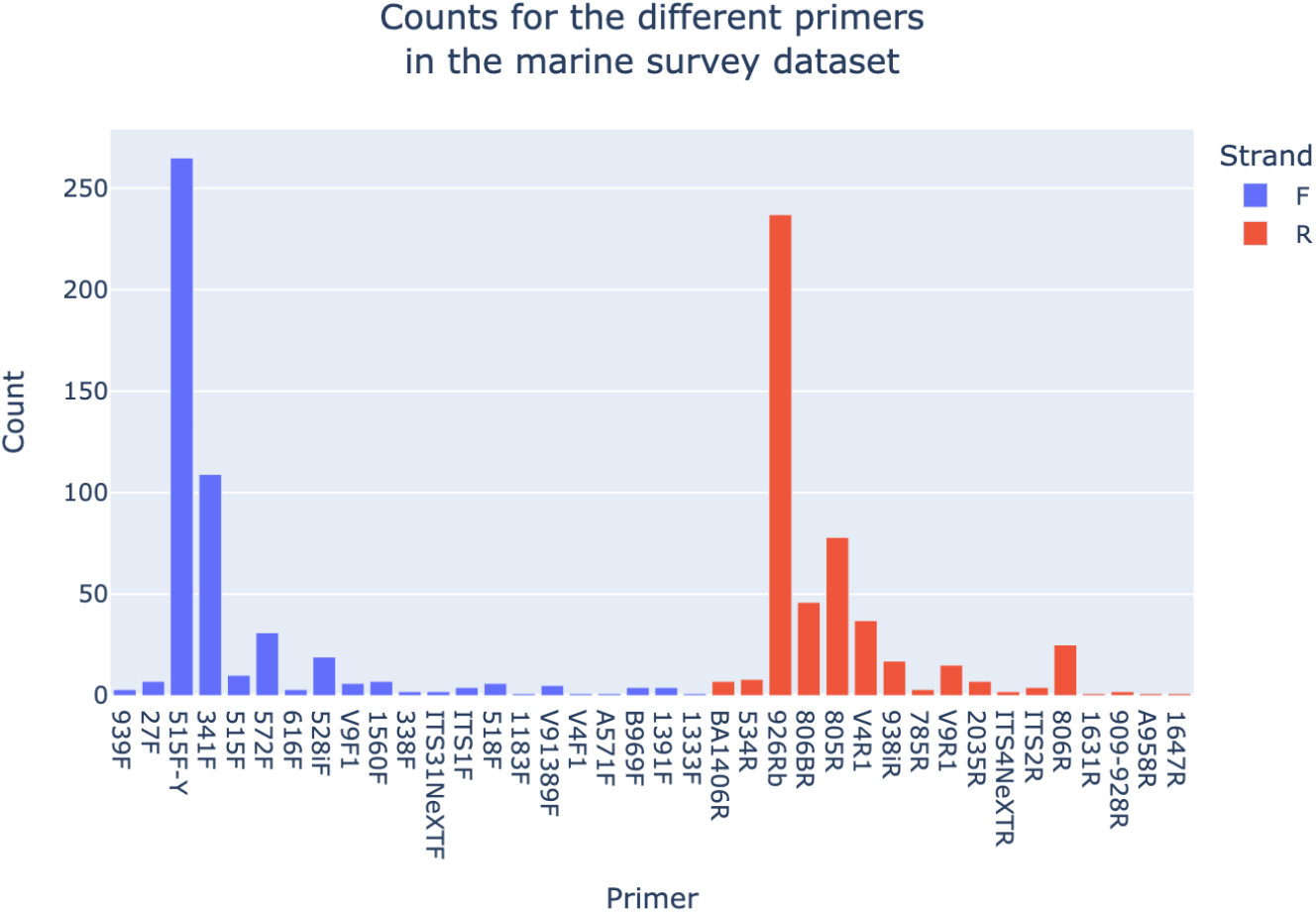
Distribution of identified standard primers in the Marine Survey Dataset for both 5^′^ to 3^′^ (F) and 3^′^ to 5^′^ (R) directions. As the most common amplified region was 16S (Figure 4), the most common standard primers were also for 16S.

### 3.3 Primer cutoff prediction inferences and primer validation strategies

The *primer cutoff prediction* inference method was developed as a secondary strategy for cases where primers are not present in the standard primer library bundled with PIMENTO. While it is less reliable than the *standard primer* method due to its heuristic nature, it is applicable to any type of metabarcoding studies, and to any type of sequenced amplicon, from rRNA to COI. When a primer is inferred in this way, the end-product is a consensus sequence from the beginning of reads to some chosen optimal cutoff point, and will therefore naturally fit the data it was generated from.

To establish the method’s accuracy in predicting real primers, multiple examples of inferred primers from the MSD were retrieved and compared to matches found in literature, which can be seen in Table 2. In run DRR059938 shows a Large Subunit (LSU) primer pair - LSUCop-D2 - which has been used in marine metagenomics analysis using the D2 region of the LSU as a marker**[18]**. Both the forward and reverse primers map to the native sequences with exact identity and coverage, with the inferred reverse primer being slightly longer by two bases.

**Table 2:**
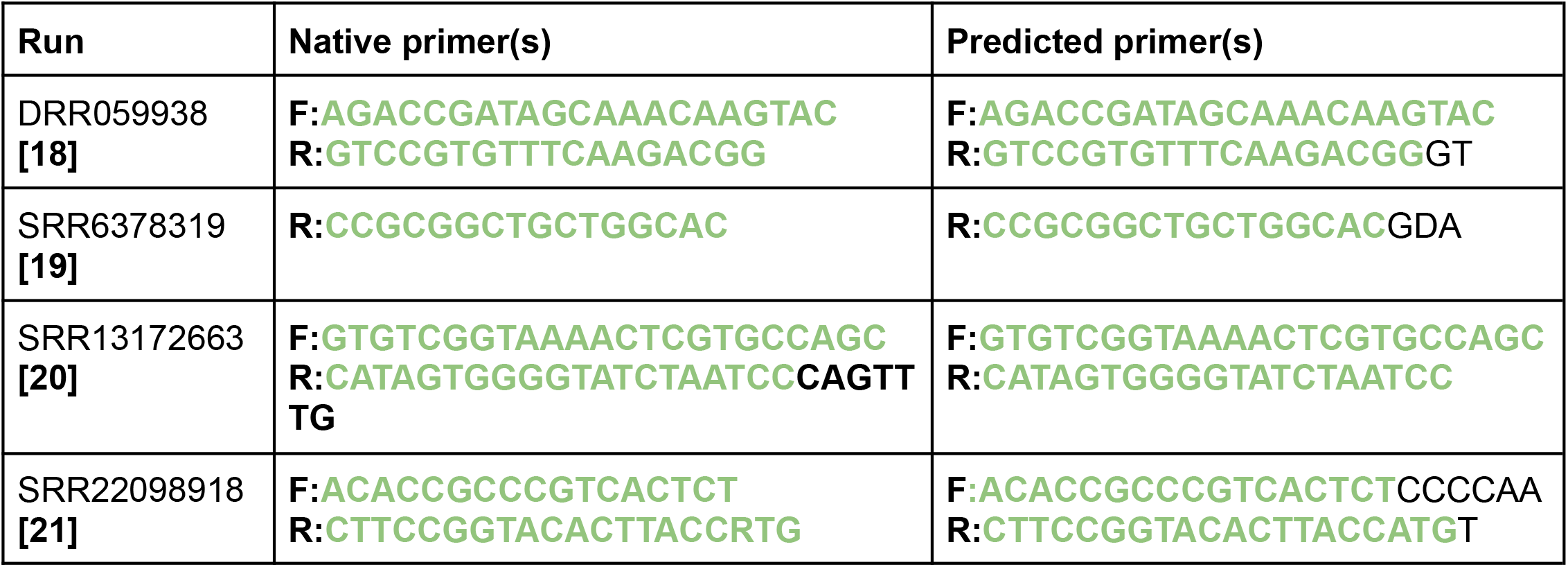
Comparison of four primer sequences that were predicted by the primer cutoff prediction method and its native counterparts, found in literature. The overlap between the native and predicted primer sequences are coloured in green. All four examples show a high coverage, and exact sequence identity for said coverage.

In run SRR6378319, a standard primer for the 16S V3 region was found that was not originally in the curated library - BV3r**[19]**. In this run, the forward primer was an existing standard primer for the V3 region: 341F. This is an example of an outcome where a standard primer was found in just one direction, but primers were predicted to exist in both directions (i.e. SsGG). In this case a standard primer was simply missing from the library, and has since been added to it. The predicted primer overlaps with 100% identity and coverage, although it does over extend by three bases.

In run SRR13172663, a primer pair for marine eDNA studies**[20]** was found in run SRR13172663. This pair is actually a universal set for fish eDNA metabarcoding, with the forward primer matching perfectly. The reverse primer is however missing 7 bases out of 20, which is explained by the cutoff prediction strategy preferring to select shorter primers as part of its algorithm, as to avoid trimming biological bases and losing potential real diversity in the process. Finally, in run SRR22098918, a different marine eDNA primer pair was found, which is often compared to the MiFish universal set**[21]**. Sequence identity was again exact, with a small number (6 and 1) of extended bases for both forward and reverse directions.

These results show that the *primer cutoff prediction* strategy for inference is marker-agnostic. While the MSD was created by retrieving sequencing runs that were annotated as originating from microbial metabarcoding experiments, incorrect annotations in public databases resulted in some eDNA studies being selected, for which primers were still able to be inferred. A further aim was to identify sequences that could be considered for addition to the standard library, such as the V3 BV3r primer. It is encouraging to note that on every occasion demonstrated in these results, the primers matched with exact sequence identity to the native primers, with no gaps or mismatches. Nonetheless, the inferred primers are not perfect: some are longer than the native primer, which could result in removal of biological diversity, and some are shorter, which could still impact ASV calling results as the primer is not fully removed. Also, there are likely to be false positives, as some runs will have been flagged for primer inference when unnecessary.

### 3.4 Consistency across biomes

The SSD and HGSD datasets were built and analysed by PIMENTO to assess the consistency of performance across different biomes. In Figures 6 and 7, the distribution of outcomes of the *standard primer search* strategy and the optional preliminary step of the *primer cutoff prediction* strategy for the SSD and HGSD can be seen, respectively, using the same interpretation of high and low confidence outcomes as used in section 3.1.

**Figure 6:**
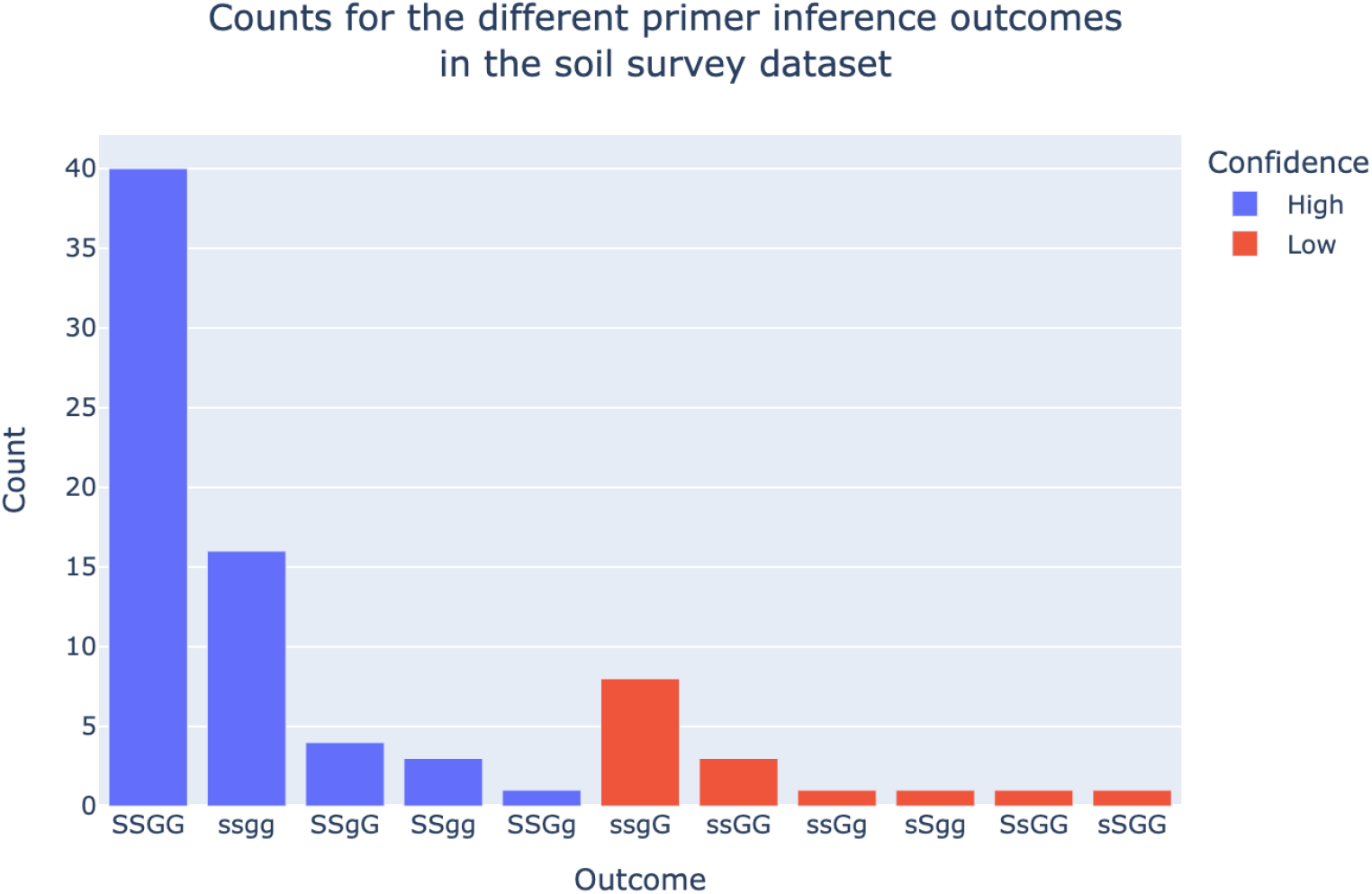
Bar plot showing the distribution of primer inference outcomes for the 79 runs of the Soil Survey Dataset. The outcomes are classified into high, and low confidence as before, with 81.0% of the outcomes considered high confidence.

**Figure 7:**
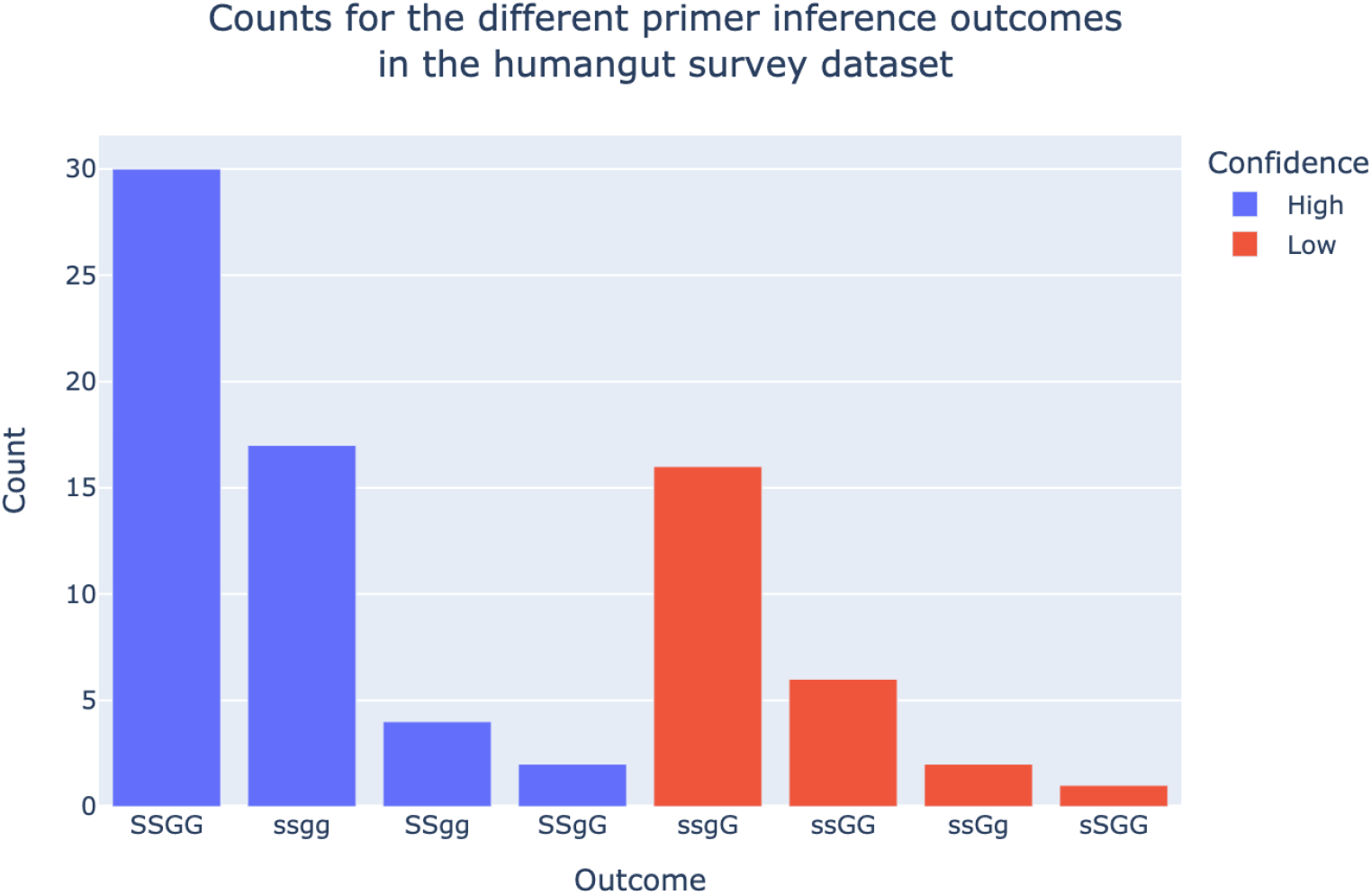
Bar plot showing the distribution of primer inference outcomes for the 78 runs of the Human Gut Survey Dataset. The outcomes are classified into high and low confidence as before, with 67.9% of the outcomes considered high confidence.

Of the 79 soil runs, 64 were assessed to be high confidence outcomes (81.0%) and 15 were low confidence outcomes (18.9%). These results are very similar as those of the marine dataset, showing consistency across these two biomes. Meanwhile, 53 of the 78 human gut runs were to be high confidence outcomes (67.9%) and 25 were low confidence outcomes (32%). While these results are still mostly made up of high confidence outcomes, the low confidence outcomes are approximately twice as common in the human gut dataset compared to both the marine and soil datasets. A deeper investigation of these low confidence cases confirmed they are runs for which primers were already removed, but with highly conserved bases at the beginning of reads, leading to false positive primer predictions.

### 3.5 Primer validation of automatically-inferred primers

As was discussed in section 3.4, one potential source of error in PIMENTO is the false positive inference of primers existing in reads when it is not the case. To prevent such false positives, validating low confidence automatically-inferred primers is always recommended. This validation can be performed either by referring to existing literature that describes the primer, or by computational means. For example, using a tool like Infernal/cmsearch with the different Rfam covariance rRNA models as input e.g. RF00177 for the bacterial 16S region, one could make sure that the generated primer sequence lies within coordinates that we would expect for a primer, i.e. before or after a hypervariable region, rather than within one.

## 4. Conclusion

In this work, we have introduced PIMENTO, which is a set of Python scripts that implement a novel dual-strategy method for filling an important gap in large-scale ASV analysis of metabarcoding data. Using either a curated library of standard primers to search against, or base-conservation patterns to automatically predict good cutoff points for inferring a consensus sequence, we believe this toolkit will help enable analysis pipelines that generate ASVs at scale.

As was shown in section 3.3, the automatically inferred primers generated by the cutoff-based method have exact sequence identity to native primers, but can sometimes be either slightly shorter or longer than the correct sequence. During the course of this work, multiple primers that were not originally in the standard primer library, but were predicted as existing and then generated by the primer cutoff prediction method, were subsequently added to the standard primer library, including the BV3r primer that was discussed in section 3.3. The addition of further primers to the library when they are first encountered means that future runs using them will benefit, as they will now be detected by the standard primer strategy directly, leading to more precise primer trimming. There is therefore significant potential in the crowdsourcing of further additions to the standard primer library, e.g. through the creation of new issues on PIMENTO’s GitHub repository, naturally improving PIMENTO’s detection capabilities over time.

## Source code, data, and package availability

The source code for PIMENTO is publicly available on GitHub at this link: https://github.com/EBI-Metagenomics/PIMENTO. The files making up the standard primer library can be found in the repository at this link: https://github.com/EBI-Metagenomics/PIMENTO/tree/main/pimento/standard_primers. Finally, PIMENTO is available on both PyPi and bioconda as the “mi-pimento” package.

